# Gene-Specific Methylation Profiles for Integrative Methylation-Expression Analysis in Cancer Research

**DOI:** 10.1101/618033

**Authors:** Yusha Liu, Keith A. Baggerly, Elias Orouji, Ganiraju Manyam, Huiqin Chen, Michael Lam, Jennifer S. Davis, Michael S. Lee, Bradley M. Broom, David G. Menter, Kunal Rai, Scott Kopetz, Jeffrey S. Morris

## Abstract

DNA methylation is a key epigenetic factor regulating gene expression. While promoter-associated methylation has been extensively studied, recent publications have revealed that functionally important methylation also occurs in intergenic and distal regions, and varies across genes and tissue types. Given the growing importance of inter-platform integrative genomic analyses, there is an urgent need to develop methods to construct gene-level methylation summaries that account for the potentially complex relationships between methylation and expression. We introduce a novel sequential penalized regression approach to construct gene-specific methylation profiles (GSMPs) which find for each gene and tissue type a sparse set of CpGs best explaining gene expression and weights indicating direction and strength of association. Using TCGA and MD Anderson colorectal cohorts to build and validate our models, we demonstrate our strategy better explains expression variability than standard approaches and produces gene-level scores showing key methylation differences across recently discovered colorectal cancer subtypes. We share an R Shiny app that presents GSMP results for colorectal, breast, and pancreatic cancer with plans to extend it to all TCGA cancer types. Our approach yields tissue-specific, gene-specific sparse lists of functionally important CpGs that can be used to construct gene-level methylation scores that are maximally correlated with gene expression for use in integrative models, and produce a tissue-specific summary of which genes appear to be strongly regulated by methylation. Our results introduce an important resource to the biomedical community for integrative genomics analyses involving DNA methylation.

## Introduction

DNA methylation involves the addition of a methyl group to cytosine residues, predominantly in the context of CpG dinucleotides in a DNA sequence. It is among the best studied epigenetic modifications, plays an important role in the regulation of gene transcription, and is associated with numerous key biological processes and diseases. For example, hypomethylation of tumor enhancer genes and hypermethylation of tumor suppressor genes have been implicated as key factors in many cancers.

With the increasing availability of multi-platform genomics and epigenomics datasets in cancer and other diseases, many researchers are interested in performing integrative analyses relating the various platforms to each other. Integrative analyses that take into account the interplatform relationships suggested by the underlying biology have the potential to help prioritize discoveries that are most likely to be functionally relevant, or even reveal insights that would be missed by analyses that only look at one platform at a time. For example, many methylation studies have focused on the identification of tissue-specific or cancer-specific differentially methylated regions. Compared to a CpG site that is differentially methylated across tissue types but not associated with gene expression, focusing on differentially methylated CpGs whose methylation is strongly correlated with expression might lead to more meaningful results. Therefore, identifying a list of CpGs associated with expression of a given gene would be helpful in guiding which CpGs to focus in methylation studies.

Some researchers have developed unified modeling frameworks for integrative analyses such as iBAG (integrative Bayesian Analysis of Genomics Data) (Wang et al. 2012; Jennings et al. 2013) and iCluster (integrative clustering) (Shen et al. 2009). iBAG is a hierarchical modeling framework that decomposes gene expression into components explained by various upstream genetic and epigenetic platforms such as copy number, methylation, and miRNA, and then incorporates these components as predictors in a clinical model. This model identifies which genes and pathways have expression levels related to clinical outcomes, while simultaneously identifying which upstream platforms appear to be driving the association. iCluster fits a joint latent variable model for integrative clustering of data based on multiple genomic platforms, including copy number, methylation, and gene expression. Integrative models like iBAG and iCluster require calculation of gene-level summaries for each genomic platform. For a platform such as copy number, it is relatively easy to come up with a reasonable strategy for computing gene level summaries (e.g. average copy number in coding region of gene), but a simple strategy like this may not work well for platforms like methylation that affect expression in more complex and subtle ways. In existing literature, we have encountered two strategies for constructing gene-level methylation summaries: (1) computing the average methylation level across all probes located in the gene’s promoter region, or (2) using the methylation level for the single probe that appears to be most negatively correlated with gene expression. While reasonable, both of these strategies appear to be simplistic and could miss the most important epigenetic effects for a given gene.

This can be seen in a study of methylation and expression of the genes EREG and AREG in colorectal cancer (CRC) that motivated this work (Lee et al. 2016). It was hypothesized that (a) higher gene expression of EREG and AREG, which encode EGFR ligands epiregulin and amphiregulin, is associated with increased sensitivity to anti-EGFR therapy, and (b) this expression is largely modulated by methylation. Initially, their integrative analysis of methylation and expression focused on the single CpG site within the promoter region of each gene that was most negatively correlated with gene expression using a pan-cancer analysis spanning all cancers in The Cancer Genome Atlas (TCGA). Fitting the iBAG model to this data, ~ 57% – 65% of expression variability was explained by methylation for EREG, while for AREG, surprisingly less than 10% of gene expression was found to be explained by methylation in the CRC samples. Upon further examination, cg03244277 was the CpG site whose methylation was most correlated with expression when looking across all cancers, but for CRC the methylation at this site had very little association with AREG expression. However, the methylation for two other sites, cg02334660 and cg26611070, had strong association with AREG expression (34% and 35% of expression variability explained, respectively), and one of these markers was located in the gene body of AREG, not the promoter region. This demonstrated that (1) functionally important methylation can vary across cancer types, (2) not all important methylation occurs in the promoter region of the gene, and (3) the choice of any single CpG to represent methylation level for a given gene has the potential to miss important signals. Together, these suggest a more comprehensive approach looking at multiple CpG sites is warranted. If the original CpG derived from the pan-cancer analysis was utilized, it would have resulted in an incorrect conclusion that AREG, a clinically important gene for CRC, is not strongly regulated by methylation. This motivated the development of a more comprehensive strategy for studying associations of methylation and expression in a tissue-specific manner, not restricting focus to promoter regions, and allowing for the potential of multiple important CpG sites per gene, to ensure that important discoveries are not missed.

Initial studies of methylation focused primarily on promoter-associated CpG islands, which are CpG-rich short regions found within the promoter regions of 70% human genes (Laurent et al. 2010) that are frequently linked with gene silencing. However, with the advent of high-throughput DNA methylation profiling methods that survey a more extensive set of CpG sites, it has become clear that the relationship between DNA methylation and gene transcription is much more complicated than previously expected (Teschendorff and Relton 2018). For instance, the comprehensive high-throughput array-based relative methylation analysis of human colon cancer methylome showed that methylation in CpG shores, flanking regions up to 2kb away from CpG islands, is strongly related to gene expression (Irizarry et al. 2009). Extensive positive correlations between gene body methylation and gene expression have been observed and reported in multiple genome-wide studies of epigenomic and gene expression data (Kulis et al. 2013). Moreover, recent studies have suggested regulatory roles for CpGs in distal elements such as enhancers. A joint analysis of the relationship among DNA methylation, sequence variation and expression variation from 62 primary fibroblast samples has found that among methylation-correlated genes, more than one third show only correlation with distal CpG sites (Wagner et al. 2014). A number of genome-wide correlation-based studies have demonstrated a negative correlation between enhancer methylation and expression of the target gene (Kulis et al. 2013). In particular, in a systematic analysis of distal methylation sites associated with gene expression in 58 human cell types, Aran et al. (2013) concluded that enhancer methylation is even more predictive of gene expression than promoter methylation. In fact, the negative correlation between enhancer methylation and target gene expression was exploited to infer the putative gene targets of active enhancers and build tumor-specific enhancer-gene networks (Yao et al. 2015; Rhie et al. 2016). In summary, these recent studies indicate that regulatory methylation sites do not occur exclusively in CpG islands or gene promoters, and the effect of DNA methylation on gene expression can depend strongly on the genomic location of the CpG site with respect to the gene region (Thingholm et al. 2016).

One simple approach to study the association of methylation and gene expression is to look at CpG sites one at a time, e.g. by calculating Spearman or Pearson correlations between gene expression and the methylation values of each CpG site in some neighborhood of the gene, ideally expanding to the flanking regions outside the gene body by at least several hundred kilobases to capture potential distal regulatory elements (Aran et al. 2013). As shown in Figure 1, for the Illumina 450k array, there are often hundreds to thousands of CpGs inside or within ±500kb flanking a gene. While this simple approach can reveal important insights, it might fail to detect some of the true associations because the surveying of such a large number CpGs per gene requires adjustment for multiple testing, and one-at-a-time analyses ignore the fact that multiple methylation sites can work together to regulate gene transcription in a coordinated manner rather than independently (Denis and Tadesse 2015). Furthermore, given that nearby CpGs tend to be correlated with each other (Leek et al. 2010), in most cases a one-CpG-at-a-time analysis will find a large number of CpGs whose methylation is associated with expression, but not all are equally important, and many are superfluous. It is likely that a small subset of these CpGs have methylation that is functionally related to expression, and the others appear to be related to expression only because of their correlation with the functionally relevant CpGs. This suggests that it may be beneficial to look at multiple CpG sites together, to identify a relatively small subset of CpGs most likely to be functionally important.

**Figure 1:**
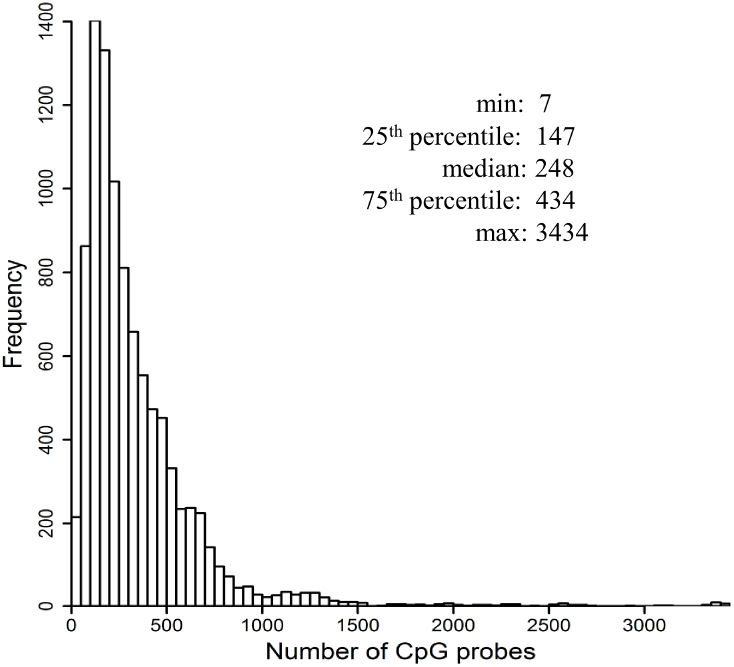
Distribution of the number of associated CpG probes per gene. For the 9,569 genes satisfying our selection criteria, the distribution of the number of CpG probes located within the gene body or the flanking region of ±500kb on either end from the Illumina 450K methylation array per gene is shown.

Under the assumption that functional importance of a particular CpG is often related to its relative location within or around the gene (Thingholm et al. 2016), several studies (Brenet et al. 2011; Lou et al. 2014; Rhee et al. 2013; VanderKraats et al. 2013; Schlosberg et al. 2017) have investigated the combinatorial effect of methylation of different components of a transcription unit on gene expression, and constructed quantitative models to predict gene expression based on methylation of different genomic regions. Among these studies, Brenet et al. (2011) concluded that compared to transcription start site (TSS), methylation in the first exon of a gene is much more tightly associated with gene silencing. Lou et al. (2014) found that while promoter and gene-body methylation are both indicative of gene expression, the latter tends to have a stronger predictive effect, and combining the methylation status in both regions leads to more accurate prediction of gene expression. With methylation data measured at single-base resolution genome-wide, such as whole-genome bisulfite sequencing, VanderKraats et al. (2013) and Schlosberg et al. (2017) modeled the spatial patterns of differential methylation within a fixed window around the TSS to predict expression changes, and found that gene-body methylation cannot further improve predictive performance relative to the promoter methylation alone.

These studies look at general quantitative relationships between methylation of different genomic regions and gene expression across the genome. However, such relationships may not apply to all genes, and specific genes may have particular methylation characteristics, suggesting that gene-specific analyses of the relationship between gene expression and methylation may be warranted. In addition, these studies do not provide gene-specific lists of which CpGs are most important, nor do they provide gene-level methylation summaries that could be used for integrative analyses. They also ignore the fact that methylation patterns tend to vary significantly across tissue types (Jones 2012), suggesting integrative analyses involving methylation be done on a tissue-specific basis. Moreover, these methods limit their analyses to CpGs in the gene body and very close to the gene body (1-2kb), whereas more distal CpGs, e.g. in enhancer regions, can also have an important effect on expression. It is also worth noting that methylation plays a fundamental role in regulating the expression levels for some genes, while it is a much less important factor for others. These general modeling approaches do not provide a summary of which genes are strongly regulated by CpG methylation.

To address these issues, we have developed a penalized regression approach to generate gene-specific methylation profiles (GSMPs) that consist of sparse lists of CpGs whose methylation levels best predict expression for each gene, and corresponding weights that indicate the relative importance and direction of their associations. Our approach considers all CpGs within the gene body or within ±500kb of the gene region, but utilizes a sequential modeling strategy that incorporates biological information from a global genome-wide analysis performed to determine which CpGs are *a priori* most likely to be associated with expression based on their characteristics. Our model first selects among the CpGs most likely to be functionally important based on their characteristics, and then considers other CpGs afterwards. This strategy is likely to yield sparser sets of CpGs that are also more likely to be functionally related to expression. GSMPs can be used to construct gene-specific methylation scores (GSMSs) that are maximally correlated with gene expression and provide a gene-specific measure of percentage of expression variability explained by methylation status of that gene.

We first develop the GSMPs in the setting of CRC, using TCGA tumor samples to build the models, and assessing the performance in terms of sparsity and ability to explain expression variability using cross validation as well as an independent validation data set consisting of tumor samples from The University of Texas M.D. Anderson Cancer Center (MDACC). We also apply this approach to produce GSMPs for breast cancer (BRCA) and pancreatic cancer (PAAD). GSMPs are constructed separately for each cancer since methylation patterns vary strongly across tissue types (Jones 2012), and as illustrated by our AREG/EREG example, global modeling across all tissue types may miss out on relationships specific to a particular tissue type. We develop a freely available Shiny App containing the GSMPs of all genes in the genome for each of the 3 tumor types.

## Results

### Selection probability functions

Figure 2 shows the results of our genome-wide analysis of CpG selection probability in terms of importance for predicting gene expression as a function of its relative location to TSS and TES (transcription end site), CpG type and gene size, for methylation sites positively or negatively correlated with gene expression.

**Figure 2:**
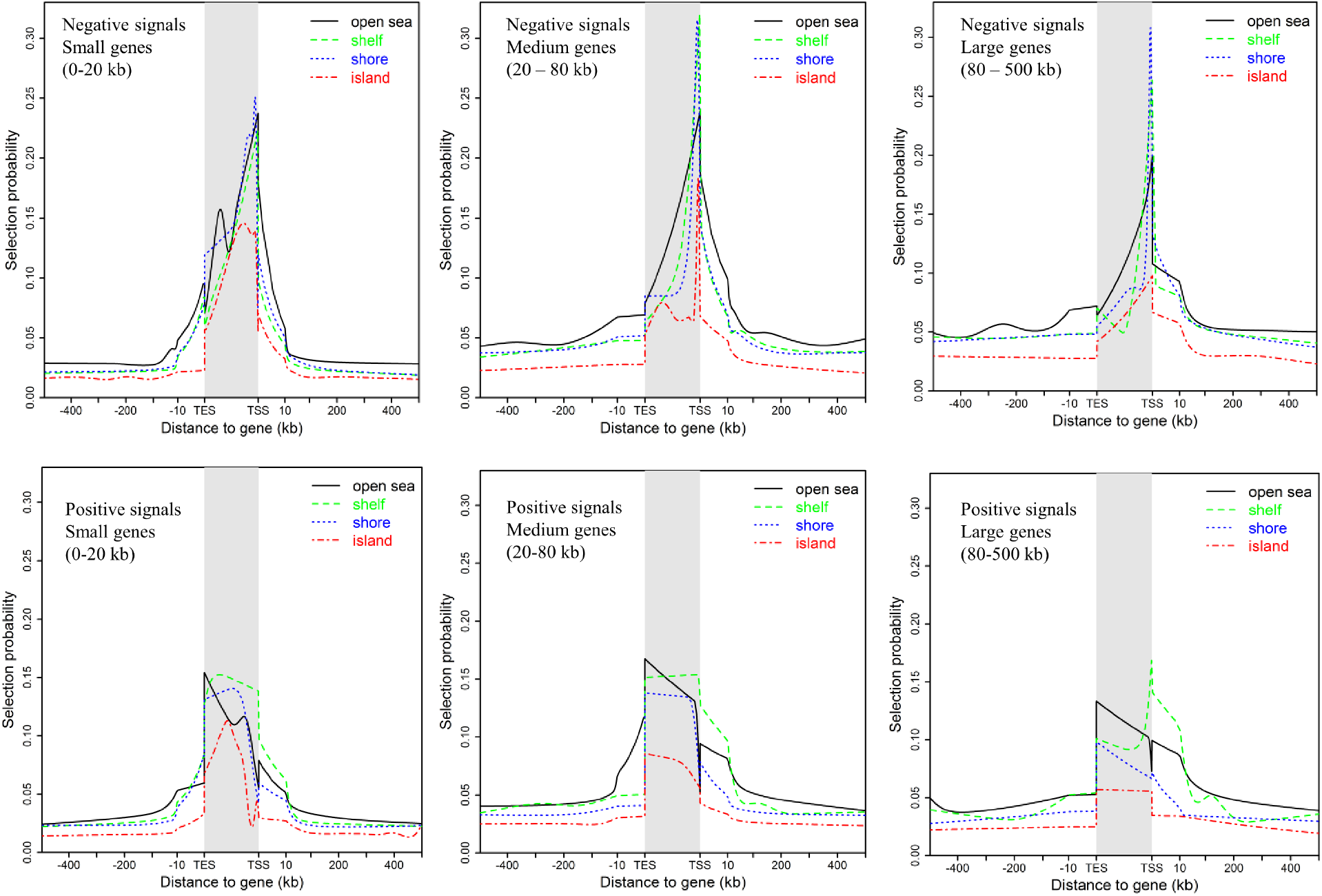
The probability of selection as a function of a CpG’s epigenomic characteristics. Each subfigure shows the probability that a CpG probe is selected (indicated by *y*-axis) as a function of its relative distance to the TSS and TES of the associated gene (indicated by *x*-axis), CpG type (CpG island, CpG shore, CpG shelf, open sea), gene size (small, medium, large), separately for CpGs negatively (top row) or positively (bottom row) associated with gene expression.

The selection probability curves in Figure 2 reveal interesting patterns. First, selection proba-bilies are always much higher for methylation sites within or very close to the gene body compared to more distant regions, where the selection probability decreases monotonically and then reaches a plateau. This suggests that functionally related CpG sites tend to concentrate around the gene body, as expected. Second, a sharp peak occurs near the TSS in the probability curve for CpG sites whose methylation levels are negatively associated with gene expression for all CpG types and gene sizes. Again, this is not surprising, as one expects that methylation in or near the promoter region of a gene should be associated with gene silencing. In contrast, the selection probability curves for positively associated CpG sites do not show this pattern, but instead show higher probabilities across the gene body, in most cases highest near the TES. Third, when controlling for direction of association and gene size, differences between the selection probability curves across CpG types are evident. Methylation sites in CpG islands are least likely to be selected in all cases.

We formally tested if CpG sites are significantly more likely to be chosen for certain CpG types and locations. Preliminary analyses suggested that the relationships among the CpG types varied across positively and negatively correlated CpG sites and inside/outside the gene body, but not by more specific location or gene size, so for simplicity we report analyses aggregating over these factors. The odds ratios of selection for each pair of CpG types along with the corresponding p-values are shown in Figure 3.

**Figure 3:**
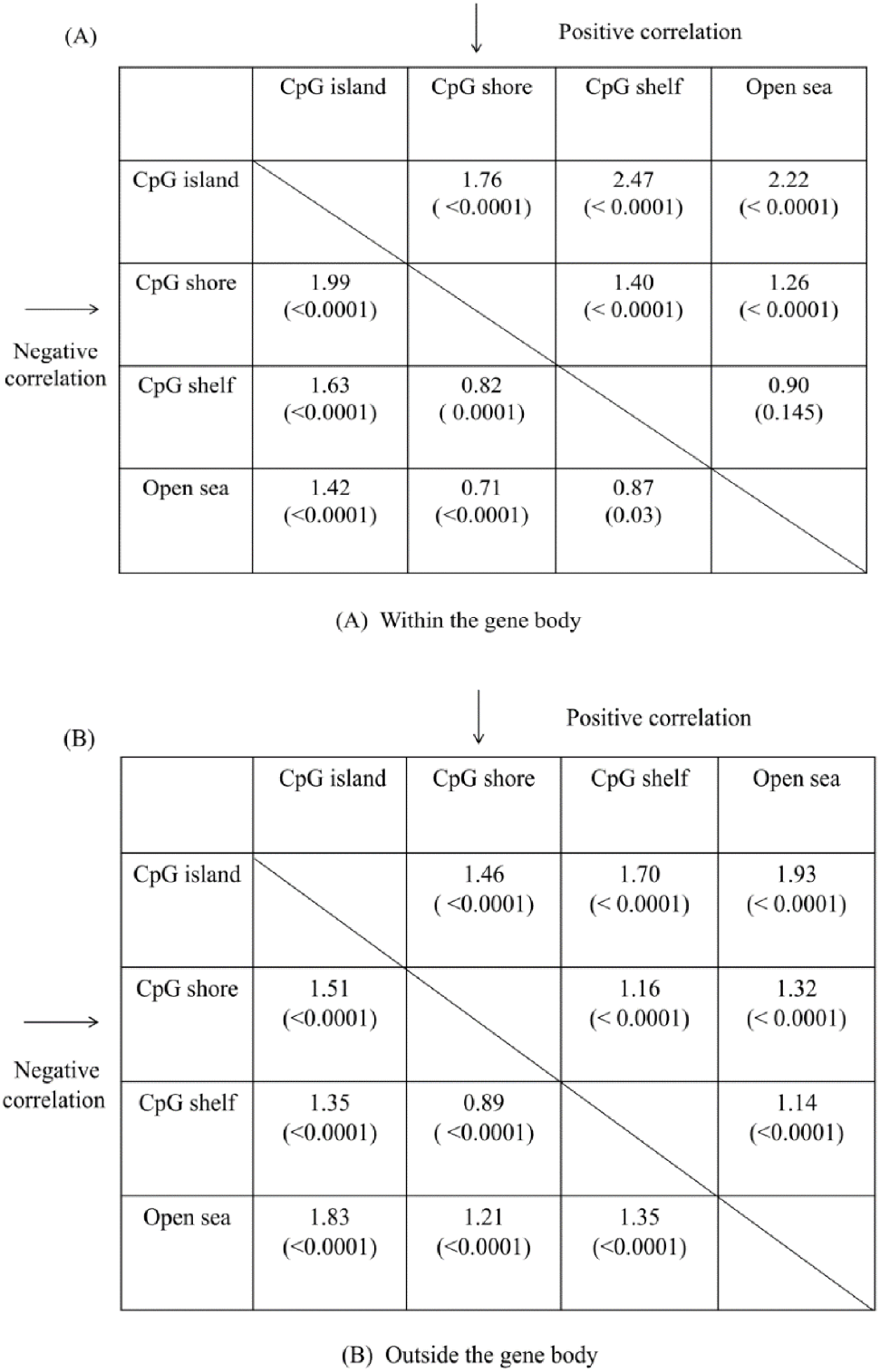
The odds ratios of selection for each pair of CpG types. For every pair of CpG types, the odds ratios of selection are shown for CpGs (A) inside the gene body and (B) outside the gene body. In each table, the odds ratio of every pair of CpG types (CpG type in the column header as numerator; CpG type in the row header as denominator) for positively correlated CpGs are shown in the upper half; the odds ratio of every pair of CpG types (CpG type in the row header as numerator; CpG type in the column header as denominator) for negatively correlated CpGs are shown in the lower half. For example, for a positively correlated CpG in the gene body, a CpG located in a shelf is 40% more likely to be selected than one from a CpG shore (0R=1.40, p≤1e-4).; for negatively correlated CpG sites in the gene body, a CpG located in the shelf is 18% less likely to be selected than one from a CpG shore (0R=0.82, p=1e-4).

For positively correlated CpG sites, the ranking of selection probabilities in decreasing order is CpG shelf ≈ open sea > CpG shore > CpG island for locations inside the gene, and open sea > CpG shelf > CpG shore > CpG island for locations outside the gene. For negatively correlated methylation sites, the ranking is CpG shore > CpG shelf ≈ open sea > CpG island for locations inside the gene, and open sea > CpG shore > CpG shelf > CpG island for locations outside the gene. Noted differences were statistically significant (p-value ≤ 0.0001 corrected for multiple testing). In all cases, CpG sites in CpG islands were least likely to be selected as important for predicting expression. Open seas were most likely to be selected for positively correlated CpG sites and all CpG sites outside the gene body, while CpG shores were most likely to be selected for negatively correlated CpG sites within the gene body.

### Comparison of sequential lasso with other methods

Next, we assessed the performance of our sequential lasso method (described in Methods) relative to other methods for identifying GSMPs in terms of percent of expression variability explained and model sparsity. We compared the sequential lasso with four alternative methods.

1. NegCor: Select the single CpG that is most negatively correlated with gene expression among methylation sites within the gene body or up to 2kb upstream of TSS.
2. AveProm: Use the average methylation level of CpGs located in a 2kb neighborhood centered at TSS to correlate with expression.
3. Lasso: Apply lasso regression of expression data on methylation data of CpGs located within the gene body or 500kb on either end.
4. Elastic net: Apply elastic net regression of expression data on methylation data of CpGs located within the gene body or 500kb on either end.

Among these alternatives, NegCor and AveProm are widely adopted in integrative studies to integrate methylation and gene expression, and the latter two are natural statistical learning strategies that one might try, but to our knowledge have not been used for this purpose in existing literature. For lasso and elastic net, the tuning parameters are chosen by 5-fold cross validation.

We applied all five methods to each of the 9, 569 genes satisfying the selection criteria (described in Methods), and evaluated the performance of each method using 3-fold cross validation on the TCGA cohort and independent validation on the MDACC cohort. For cross validation, we calculated the Spearman correlation between the predicted and actual gene expressions across 369 samples in the TCGA cohort. For independent validation, we first obtained the GSMP from the TCGA data and used it to compute the predicted expressions for the MDACC integromics cohort, and then calculated the Spearman correlation between the predicted and actual gene expressions. We used Spearman correlations because the MDACC integromics cohort used a different platform (Agilent) to measure gene expression than TCGA (RNAseq), leading to gene expression measurements on different scales. To focus on genes with a reasonably high percentage of expression variability explainable by methylation, for these comparisons our attention was restricted to the 4, 598 genes with a Spearman correlation of at least 0.40 between the actual gene expressions and predicted expressions based on 3-fold cross validation of the TCGA data using sequential lasso.

The distributions of Spearman correlations between actual and predicted expressions of the 4,598 genes for each method are shown in Figure 4B and 4C. Comparing the standard methods commonly used in the literature, the NegCor tends to lead to significantly higher correlations than the AveProm, but all three penalized regression approaches (lasso, sequential lasso and elastic net) have substantially better predictive performance than both of these methods, suggesting that useful information is lost by collapsing all methylation information into one single site per gene. This difference is seen in both the TCGA data and the independent validation data. Note that across all methods the Spearman correlations are lower in the independent validation data set, which is at least partially due to the different gene expression platforms.

**Figure 4:**
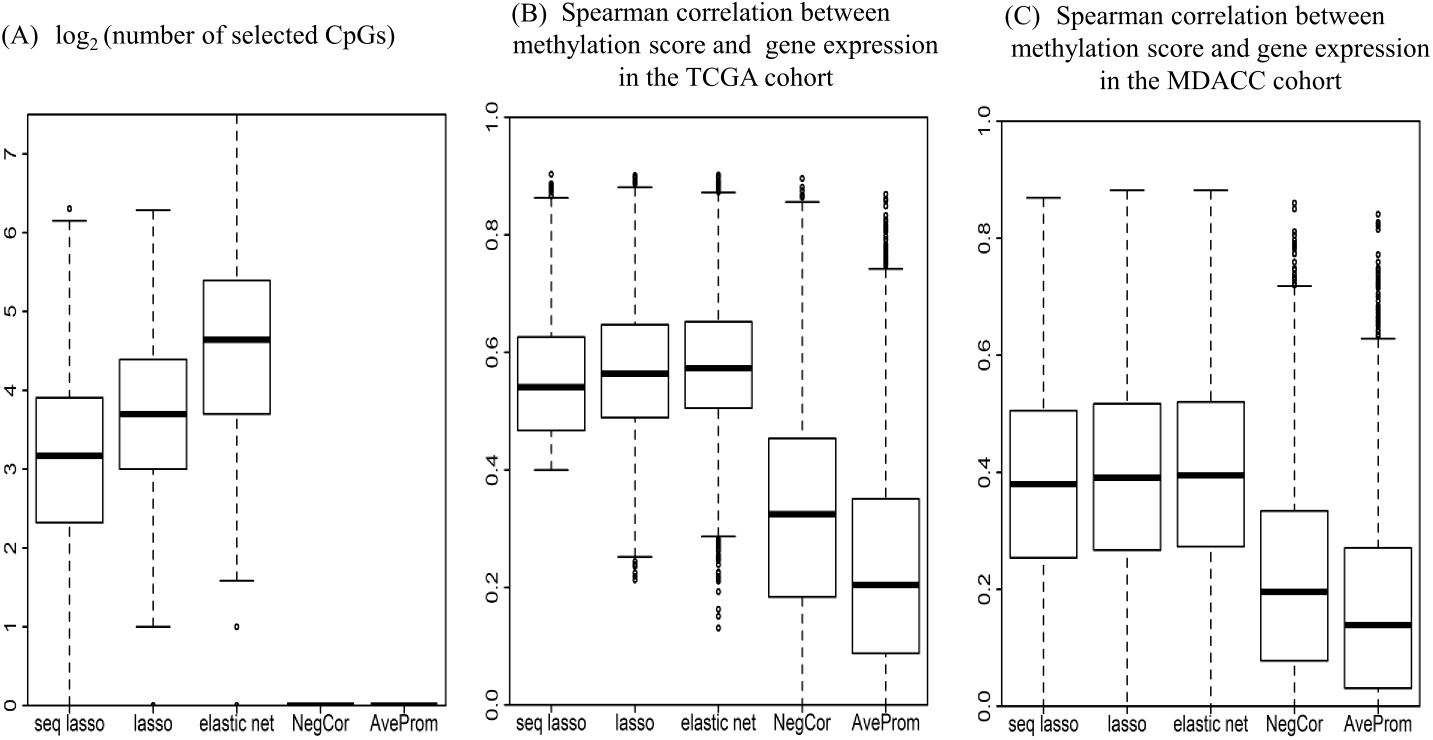
Performance comparison of the sequential lasso with other alternative methods. For the 4,598 genes with a Spearman correlation of at least 0.40 between actual and predicted gene expressions using sequential lasso in the TCGA cohort, the subfigures compare the performance of sequential lasso with lasso, elastic net, NegCor and AveProm for identifying GSMPs in terms of sparsity (A), and the ability to explain expression variability in the TCGA cohort (B) and the MDACC cohort (C).

For the 3-fold cross validation of the TCGA data, the correlations for the three penalized regression approaches are similar, with lasso and elastic net tending to have slightly higher correlations (median = 0.564 for lasso; median = 0.573 for elastic net) than sequential lasso (median = 0.541). However, this marginal improvement is achieved at the cost of incorporating many more CpGs. Figure 4A contains boxplots of the number of CpGs selected by each method. Sequential lasso selects a much smaller number of CpGs than either lasso or elastic net. The median number of CpGs selected by sequential lasso is 9, nearly two-thirds of the median for lasso (13) and one-third of the median for elastic net (25). This demonstrates that sequential lasso tends to give a more sparse and parsimonious set of important CpGs. For the independent validation, the Spearman correlation distributions for sequential lasso (median = 0.380), lasso (median = 0.391) and elastic net (median = 0.395) are very similar for the three penalized regression approaches, and again much higher than the NegCor (median=0.196) or the AveProm (median=0.139).

There are 3,248,385 gene-CpG pairs for the selected 9,569 genes. Among them, 77,626 pairs (2.4%) are selected by sequential lasso, with 34,392 (44.3% of the selected pairs) positively correlated and 43,234 (55.7% of the selected pairs) negatively correlated with gene expression. Among these 9,569 genes, there are 7,441 genes (77.8%) for which at least one CpG outside the gene body is selected by sequential lasso; 5,803 / 5,439 / 4,848 / 3,679 genes (60.6% / 56.8% / 50.7%/ 38.4%) whose GSMPs produced by sequential lasso include at least one CpG located at least 100/200/300/400kb away from the gene body. Figure S1 shows the distribution of Spearman correlations between actual gene expressions and predicted expressions in the TCGA cohort using sequential lasso across all the 9,569 genes. Based on 3-fold cross validation Spearman correlation, these 9,569 genes can be classified into 4 categories: 1) little to no association between methylation and gene expression (correlation < 0.2; 2,807 genes) 2) very weak association between methylation and gene expression (correlation 0.2 ~ 0.4; 2,164 genes) 3) moderate association between methylation and gene expression (correlation 0.4 ~ 0.6; 3,124 genes) 4) very strong association between methylation and gene expression (correlation ≥ 0.6; 1,474 genes).

## GSMPs

### Motivating example: AREG and EREG

For AREG and EREG from the motivating example of this study, the GSMPs produced by sequential lasso and other methods are shown in Figure 5A-C and 6A-C, with selected CpGs and their weights highlighted in the figure. The gene body is indicated by the dark gray bar, with an arrow demonstrating the direction of transcription, and the nearby flanking genes are indicated by light gray bars with gene names listed at the top.

**Figure 5:**
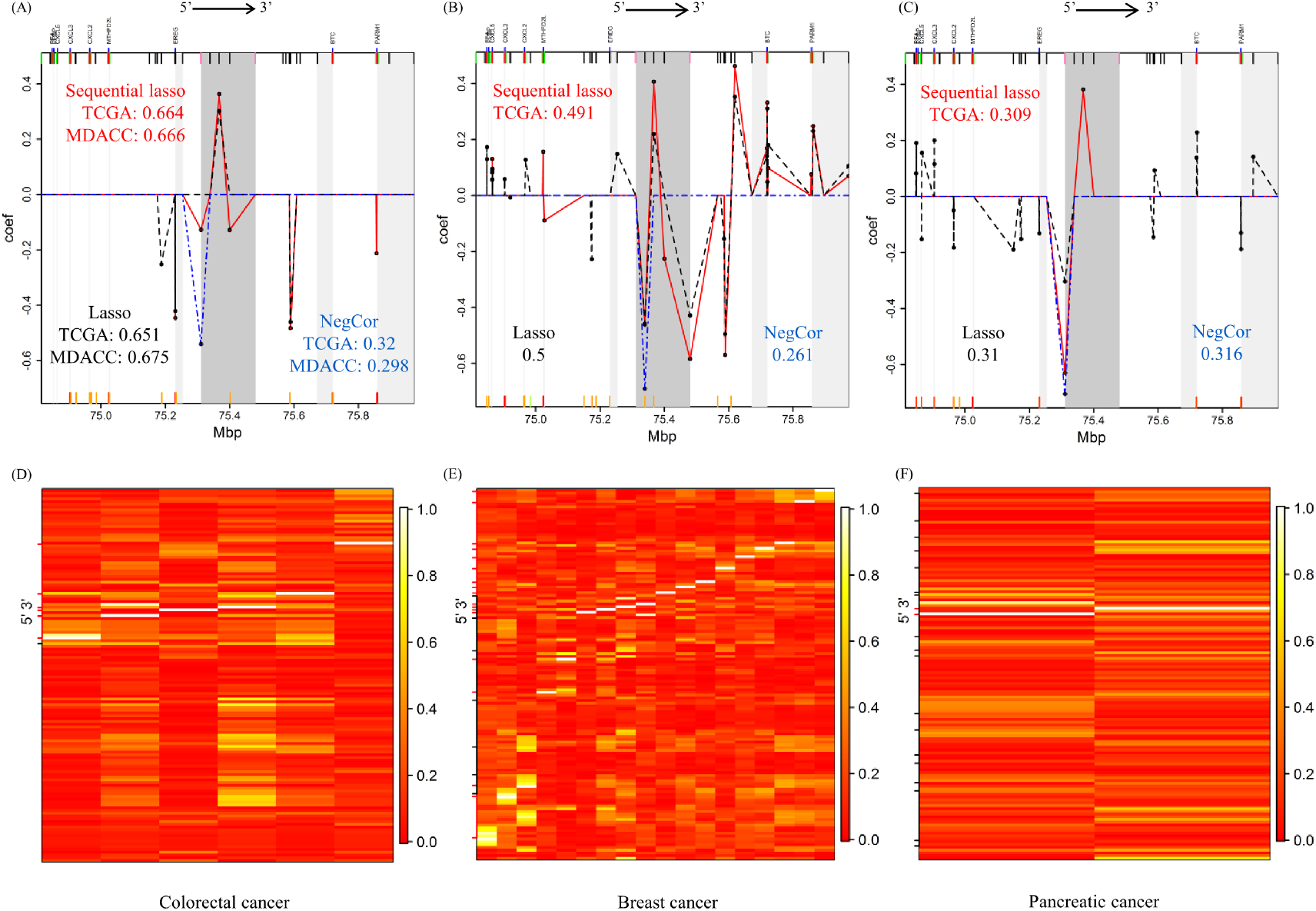
AREG methylation profile and heatmap of absolute correlations across CpGs. (A)-(C) show the GSMPs identified by different methods, with the marks at the top indicating all CpG sites in the region (CpG island-red; CpG shore-pink; CpG shelf-green; open sea-black), and the marks at the bottom indicating active chromatin states predicted using ChromHMM (active TSS-red; flanking TSS-orange red; active enhancer-orange; transcribed enhancer-green yellow). The heatmaps (D)-(F) show the pairwise absolute Pearson correlation coefficients between CpGs selected by sequential lasso (arranged in columns) and all CpGs within ±500kb of AREG (arranged in rows), with the red / black marks on the left indicating CpGs selected by sequential lasso / lasso but not sequential lasso.

**Figure 6:**
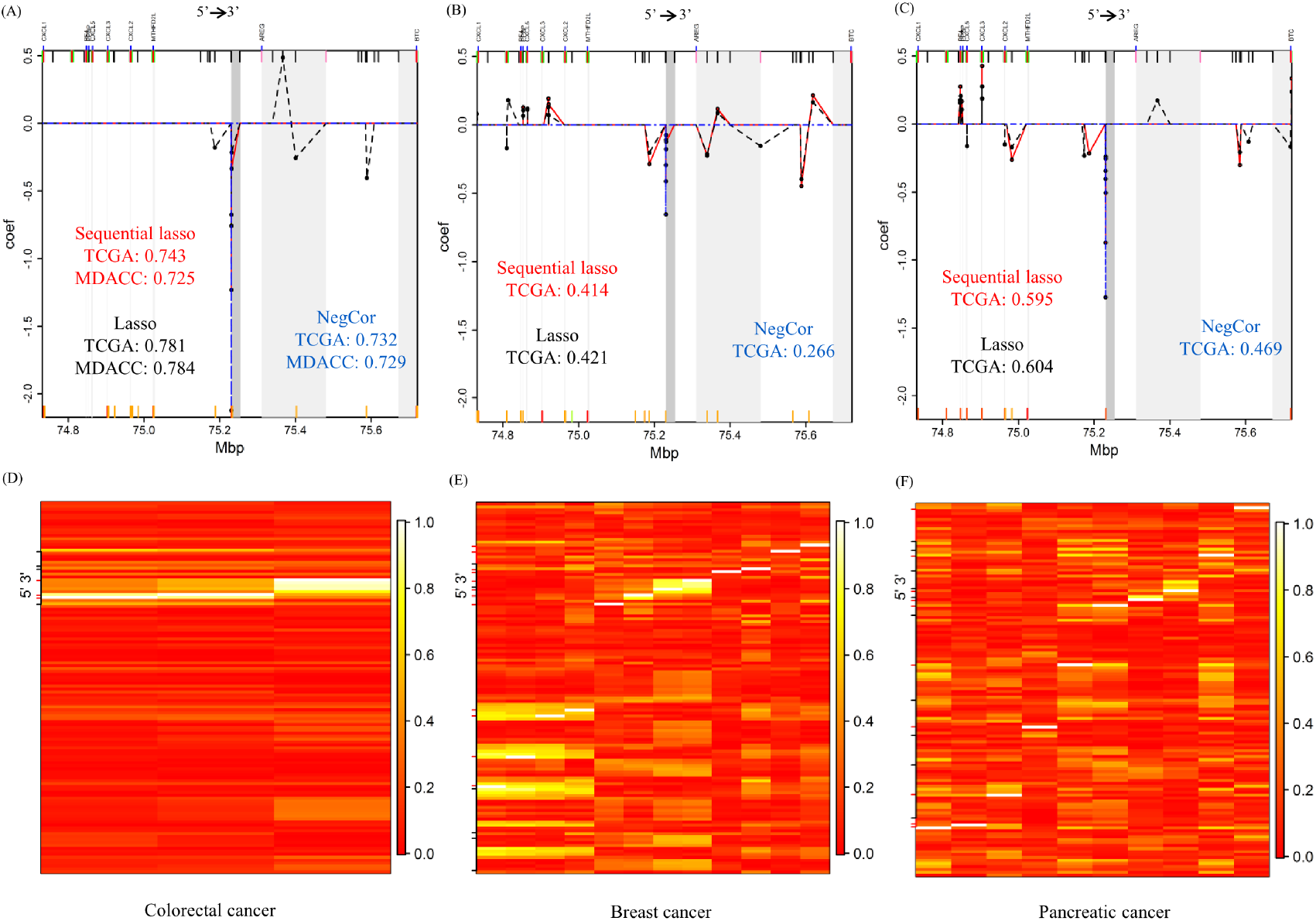
EREG methylation profile and heatmap of absolute correlations across CpGs. (A)-(C) show the GSMPs identified by different methods, with the marks at the top indicating all CpG sites in the region (CpG island-red; CpG shore-pink; CpG shelf-green; open sea-black), and the marks at the bottom indicating active chromatin states predicted using ChromHMM (active TSS-red; flanking TSS-orange red; active enhancer-orange; transcribed enhancer-green yellow). The heatmaps (D)-(F) show the pairwise absolute Pearson correlation coefficients between CpGs selected by sequential lasso (arranged in columns) and all CpGs within ±500kb of EREG (arranged in rows), with the red / black marks on the left indicating CpGs selected by sequential lasso / lasso but not sequential lasso.

Figure 5A shows the GSMP of AREG in colorectal cancer. Both sequential lasso and lasso identify a CpG probe in the middle of the gene body whose methylation exhibits strong positive correlation with gene expression, and capture two CpG sites located at a distance from the gene body of AREG, for which the methylation levels are strongly inversely related to gene expression. Interestingly, one of them is located exactly at the TSS of EREG, which is situated roughly 50kb upstream of AREG. The predicted expression by sequential lasso has a Spearman correlation of 0. 664 and 0.666 with actual AREG expression respectively for 3-fold cross-validation in the TCGA cohort and independent validation in the MDACC cohort, respectively meaning that 44.1% of AREG expression variation is explained by methylation in the TCGA cohort, and 44.4% of AREG expression variation is explained by methylation in the MDACC cohort. Both sequential lasso and lasso yield much higher Spearman correlations than NegCor (0.320 for 3-fold cross validation and 0. 298 for independent validation) and AveProm (0.333 for 3-fold cross validation and 0.298 for independent validation, not shown), indicating that these naïve methods miss functionally related CpGs when integrating methylation and gene expression.

Figure 5B and 5C show the GSMPs of AREG in breast cancer and pancreatic cancer respectively. A comparison of Figure 5A, 5B and 5C reveals that the GSMPs of AREG are very different across tissue types. Focusing on sequential lasso, the identified CpG sites have modest overlaps among three tumor types considered. In addition, the Spearman correlations between the actual and predicted expressions greatly differ across tissue types. The association between AREG expression and methylation is very strong in colorectal cancer, moderate in breast cancer, and weak in pancreatic cancer.

Figure 6A shows the GSMP of EREG in colorectal cancer. The sequential lasso identifies 3 neighboring CpG sites at the TSS of EREG as functionally important. NegCor picks out one very strong negative signal at the TSS, and yields similar Spearman correlation between the actual and predicted EREG expressions as sequential lasso, suggesting that methylation at this single CpG site can explain as much variation in EREG expression. Lasso produces a more noisy methylation profile including many more CpGs, with only marginally higher correlations.

Figure 6B and 6C show the GSMPs of EREG in breast cancer and pancreatic cancer respectively. Comparing the GSMPs in Figure 6A, 6B and 6C, the most negatively correlated CpG site remains the same across tissue types, but sequential lasso identifies some more CpGs specific to breast cancer and pancreatic cancer. Similar to AREG, the strength of association between EREG expression and methylation depends on the tissue type. EREG expression and methylation are very strongly correlated in colorectal cancer, and only moderately correlated in breast cancer and pancreatic cancer.

Figure 6D-F show the pairwise absolute Pearson correlation coefficients between the CpGs identified by sequential lasso and all the CpGs located within ±500kb of EREG, respectively for the three tumors considered. The red marks on the left of each heatmap denote the CpGs identified by sequential lasso, and the black marks denote the CpGs that are selected by lasso but not sequential lasso. In each heatmap, the rows labeled by black marks often include at least one light colored cell, indicating that a CpG identified by lasso but not sequential lasso is often highly correlated (or anti-correlated) with at least one CpG identified by sequential lasso. This phenomenon results from the sequential nature of our approach. After sequential lasso identifies important CpGs in the first step, the methylation sites that are correlated with expression because of their strong correlation with those identified CpGs would not be selected in the second step, leading to a sparser GSMP than lasso. This phenomenon can also be observed for AREG (Figure 5D-F).

### GSMP Shiny App

To facilitate the search and visualization of GSMPs, we have developed a freely available R Shiny app that can interactively display the GSMP of any gene in the genome for colorectal, breast and pancreatic cancer, which includes a tissue-specific, gene-specific list of important CpGs and their corresponding weights, along with the Spearman correlation between actual gene expression and predicted expression based on 3-fold cross validation in the TCGA cohort for each gene. Note that a measure of percent of expression variability explained by the GSMP can be computed by squaring the Spearman correlations. We show how to use the Shiny app and describe some CMS-related genes with interesting GSMP in Figure S2.

### GSMSs and consensus molecular subtypes in CRC

Guinney et al. (2015) discovered and validated four consensus molecular subtypes (CMS) of colorectal cancer, combining information across over 4000 patients from 18 studies and across 6 different subtyping systems. These subtypes appear to have different biological characteristics, with CMS1 (MSI Immune) including hypermutated and hypermethylated tumors that tend to have microsatellite instability and immune pathway activation, CMS2 (Canonical) demonstrating canonical CRC characteristics including epithelial differentiation, MYC and WNT activation, and high levels of chromosomal instability, CMS3 (Metabolic) including epithelial tumors and showing high levels of metabolic dysregulation, and CMS4 (Mesenchymal) including tumors demonstrating characteristics of epithelial-mesenchymal transition (EMT), activation of TGF-β, angiogenesis, and stromal infiltration. In spite of the fact that these subgroups were discovered based on global biology, not prognostic or predictive considerations, early results demonstrate that these CMS have potential prognostic and predictive value for precision therapy strategies. CMS4 patients have poorer prognosis and perhaps need more aggressive treatment, and they show limited benefit from Oxaliplatin (Song et al. 2016; Okita et al. 2018). Patients with CMS2 or CMS3 may benefit from addition of Bevacizumab (Mooi et al. 2018) but are not good candidates for immunotherapy (Becht et al. 2016; Lal et al. 2018), and anti-EGFR therapy appears to work very well for CMS2 patients but not CMS1 (Okita et al. 2018).

Here we compute gene level methylation-summaries (GSMS) for CRC and compare across CMS to identify genes with methylation-regulated expression levels differing across CMS. As described in Methods, GSMS can be computed from GSMP and provide a gene-level summary of functionally relevant CpG methylation activity for each subject, with high GSMS indicating that methylation profiles predict expression inhibition and low GSMS indicating that methylation profiles suggest expression enhancement.

Among the 14, 857 genes with available GSMPs, there are 7, 293 genes for which the crossvalidation Spearman correlation between actual and predicted expressions is at least 0.4 for sequential lasso, NegCor or AveProm in the TCGA cohort, suggesting moderate or strong correlation between methylation and expression in these genes. For each of the 7, 293 genes, three versions of GSMS are calculated based on GSMP produced by sequential lasso, NegCor and AveProm for each TCGA and MDACC sample, and ANOVA is performed for each version of GSMS across CMS in the TCGA cohort to detect genes whose methylation status significantly differs across CMS. At 1% false discovery rate, we detect 6,698 genes in the TCGA cohort using GSMS based on sequential lasso, greatly more than the 5,191 and 5,538 genes detected by NegCor and AveProm, respectively. The heatmaps of GSMS based on sequential lasso of the 6,698 detected genes are shown for the TCGA cohort (Figure 7A) and the MDACC cohort (Figure 7B) grouped by CMS status. The heatmaps suggest that the methylation status displays a distinctive pattern across CMS status for many genes. For example, the methylation profiles for genes located in the top thousand rows in Figure 7A tend to produce gene expression suppressed in normal or CMS1 subjects and enhanced in CMS2 subjects compared to others. Figure 7B plots the GSMS for the MDACC cohort with genes in the same order as Figure 7A, and demonstrates similar structure and independently validates these effects. The heatmaps of GSMS based on NegCor and AveProm for both cohorts are shown in Figure S3 and Figure S4, respectively. Notably, GSMS based on sequential lasso demonstrates noticeably stronger inter-CMS structure than NegCor or AveProm.

**Figure 7:**
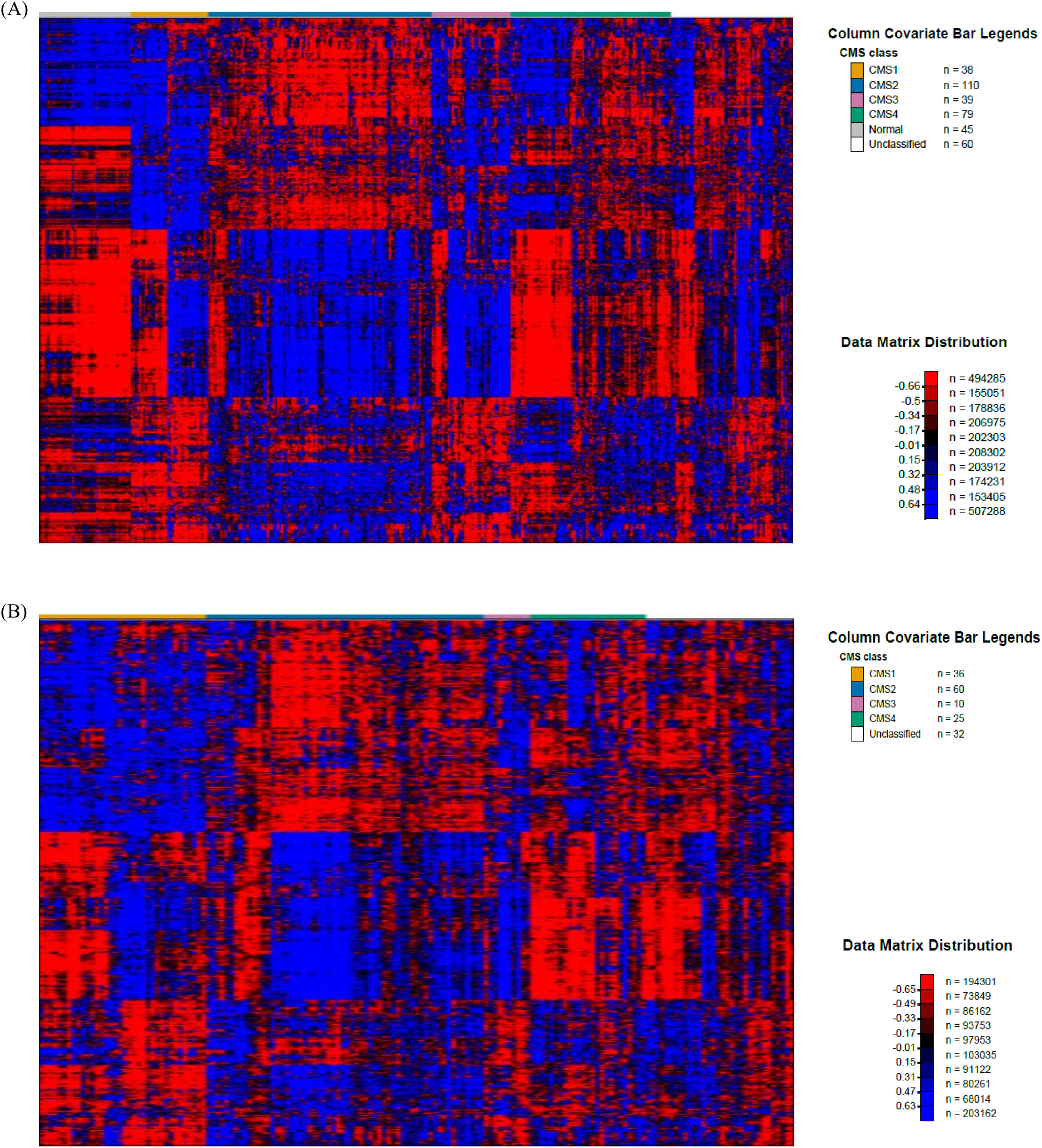
**Gene-specific methylation scores calculated using sequential lasso** of 6,698 genes whose methylation status differs significantly across CMS status in the TCGA cohort at 1% false discovery rate for A) TCGA cohort and B) MDACC cohort.

## Discussion

We propose a novel method to integrate methylation and gene expression using a sequential penalized regression approach. This approach is tissue-specific and gene-specific, and considers CpG sites within a 500kb neighborhood of the gene, where distal regulatory elements such as enhancers are usually located (Wagner et al. 2014; Aran et al. 2013). By applying an L1 penalty and using a sequential modeling approach that incorporates information from a genome-wide analysis regarding which CpGs are *a priori* most likely to be associated with expression, our approach is able to produce sparse tissue-specific, gene-specific lists of functionally important CpGs and corresponding weights, which we term as GSMPs. GSMPs can be used to construct gene-level methylation summaries that are maximally correlated with gene expressions, and produce a tissue-specific, gene-specific measure of percent variability in expression explained by methylation. A publicly available Shiny app has been developed to interactively visualize GSMP.

By incorporating prior knowledge, the sequential lasso is able to essentially accomplish the same predictive ability as straight lasso and elastic net models while using a significantly smaller set of CpGs that may also tend to be more functionally relevant and biologically interpretable. In addition, comparing our sequential lasso approach with others in CRC data demonstrates the insufficiency of summarizing gene-level methylation by taking the most negatively correlated probe or the average methylation level around the TSS, as is commonly done in practice.

The analysis of individual GSMPs reveals that the methylation profiles vary greatly across genes and tissue types, demonstrating the importance of using a tissue-specific and gene-specific approach. We also find that GSMPs often include many distal CpG sites and intergenic regions far from the promoter region, and the inclusion of these sites leads to significantly better ability to explain expression than approaches that limit attention to the gene body or promoter region.

In multi-platform integration, it is frequently useful to compute gene-level summaries of various platforms. Our approach yields gene-level summary of methylation which is sparse and maximally correlated with expression. These gene-level summaries can be used in integrative models, such as the iBAG model (Wang et al. 2012; Jennings et al. 2013) and the iCluster model (Shen et al. 2009). Gene-level methylation summaries can also be associated directly with demographical and prognostic factors, as illustrated by the ANOVA analysis of gene-specific methylation scores and consensus molecular subtypes in CRC, or clinical outcomes such as survival data. A similar strategy could be used to construct gene-level summaries for other genomic platforms, such as DNA mutation and miRNA.

To identify the CpG sites which are most likely to be associated with gene expression *a priori,* our model combined information across the genome to estimate this prior probability as a function of epigenomic characteristics of a CpG site. Apart from guiding the sequential selection, these selection probability functions constitute important biological findings by themselves, which may contribute to the unraveling of complicated regulatory roles taken by methylation. They also corroborate some previous findings regarding the methylation-expression relationship. For example, the sharp peak around the TSS observed in selection probability curves for negatively correlated CpGs lends statistical evidence to the frequently observed association between promoter hyper-methylation and gene silencing. Besides, the formal testing comparing the selection probability of different CpG types concludes that methylation sites in CpG shores are much more likely to be associated with expression among negatively correlated CpGs, which supports the finding that CpG shore methylation is strongly associated with gene expression (Irizarry et al. 2009).

It should be pointed out that our modeling approach assumed a linear regression setting for interpretability, implying the effect of methylation sites on gene expression is linear and additive (Wang et al. 2012). It would be possible to extend this approach to consider more general and flexible models with nonparametric nonlinear effects and interactions allowed, but these are outside the scope of this study. In addition, the conclusions drawn from our integrative approach should not be interpreted as strict causal relationships, but instead represent strongly associated CpG sites that are potential key methylation switches, but would need further functional validation such as modulation using CRISPR/dCas9 fusion proteins to methylate/demethylate the region to confirm any causative relationships. Finally, based on our analysis, it is clear that the use of the 27K methylation array, which contains roughly one or two CpGs per gene mostly found in the promoter region (Jeschke et al. 2015), would miss many important CpGs relative to the 450K methylation array. Recently, whole-genome bisulfite sequencing has been used to interrogate CpGs at single-base resolution (Lou et al. 2014). Given sufficient samples, we could rerun our integrative model in the future on the whole methylome data to obtain more accurate gene-level methylation summaries.

In our current Shiny R app that we will freely share upon publication, we present the GSMP results for colon, rectal, breast, and pancreatic cancer types. Our future plans include expanding results to include GSMP for all TCGA cancers, which will be included in future updates of our Shiny app.

## Methods

### Colorectal cancer data Samples

We analyzed TCGA primary solid tumor samples in colon (COAD) or rectal (READ) whose methylation data and matching gene expression data are both available, pooling them as a CRC cohort. For patients with duplicate tumor samples, one of them was randomly chosen. There are 369 samples in total. These data were used to fit our model and determine the GSMPs.

We used a cohort of 163 tumor samples from primary resection specimens from colon or rectal cancer patients at MDACC to validate our GSMPs. For these samples, both methylation data and gene expression data are available. These samples were collected between 2001 and 2009, with 87% being stage 2 and 3 cancers, and the remaining 13% stage 4. All samples had frozen tissue from primary resection specimens stored and used for expression and methylation analyses.

### DNA methylation data

We extracted DNA methylation data for TCGA samples using the R package “GeneSurvey” which can be accessed from GitHub (https://github.com/MD-Anderson-Bioinformatics/GeneSurvey); preprocessed DNA methylation data for Integromics samples were available at MDACC. DNA methylation data for both cohorts were generated from the Illumina Infinium HumanMethylation 450K BeadChip, which measures methylation level of 485, 577 CpG sites across the genome (Dedeurwaerder et al. 2011). Methylation status is quantified by beta values, the ratio of background-corrected methylated allele intensity to the sum of methylated and unmethylated allele intensities at each CpG interrogation site (Du et al. 2010). Beta values range from 0 to 1, with 0 indicating no methylation of a CpG site at either allele, 1 indicating methylation at both alleles. Possible batch effects have already been examined and corrected for both cohorts. CpG probes which can be mapped to multiple genomic locations or contain common SNPs (minor allele frequency ≥ 1%) at the CpG interrogation sites are removed, leaving approximately 392,000 CpG sites for further analysis (Chen et al. 2013).

### Gene expression data

For the TCGA cohort, we used the R package “GeneSurvey” to extract level 3 RNASeqV2 gene-level expression data, which were generated from the Illumina HiSeq platform, quantitated by RSEM (RNA-seq by Expectation Maximization), and then normalized across samples so that each sample has a fixed upper quartile value of 1000. In the “GeneSurvey” package, expression values are log-transformed, and possible batch effects have been checked and removed.

For the MDACC cohort, gene expression data were generated from Agilent microarrays. Loess based normalization was performed using the Agilent feature extraction software with background subtraction, and batch effects were checked and removed.

### Selection of genes

The generic annotation file used in TGCA RNASeqV2 analysis was downloaded from the TCGA portal (https://tcga-data.nci.nih.gov/tcgafiles/ftp_auth/distro_ftpusers/anonymous/tumor/brca/cgcc/unc.edu/illuminahiseq_rnaseqv2/rnaseqv2/unc.edu_BRCA.IlluminaHiSeq_RNASeqV2.mage-tab.1.6.0/DESCRIPTI?N.txt; accessed June 20, 2016) to obtain structure annotation information for all genes and isoforms. The genomic coordinates are based on HG19 UCSC Gene standard track (Dec 2009 version, https://tcga-data.nci.nih.gov/tcgafiles/ftp_auth/distro_ftpusers/anonymous/tumor/brca/cgcc/unc.edu/illuminahiseq_rnaseqv2/rnaseqv2/unc.edu_BRCA.IlluminaHiSeq_RNASeqV2.mage-tab.1.6.0/DESCRIPTI?N.txt; accessed June 20, 2016). There are 17, 627 genes whose expression data and methylation are both available in R package “GeneSurvey”. The genes with low inter-individual expression variation (median absolute deviation = 0) were removed from further analysis, leaving us with 14,857 genes. While we computed GSMPs for all of these genes, for model comparison and interpretation we focused on 9, 569 genes with 1) a single isoform; or 2) multiple isoforms with the same TSS and TES; or 3) at least one major isoform whose expression value is very strongly correlated with gene-level expression (Pearson correlation ≥ 0.85) across 369 TCGA samples.

### Data of other cancer types

We also applied our approach to compute GSMPs using TCGA primary solid tumor samples in pancreas or breast whose methylation data and matching gene expression data are both available. For patients with duplicate tumor samples, one of them was randomly chosen. There are 782 breast cancer samples and 178 pancreatic cancer samples. We called the R package “GeneSurvey” to extract methylation and gene expression data, for which the sequencing platforms and data preprocessing procedures are exactly the same as TCGA CRC data. We used the same generic annotation file as CRC, and also excluded genes with low inter-individual expression variation from calculation of GSMP.

### GSMP by sequential lasso Overview

Because methylation in distal regulatory elements may also be functionally related, we considered all CpGs located up to 500kb upstream of TSS through 500kb downstream of TES as potentially important. Previous studies show that distal regulatory elements are generally located within this neighborhood (Wagner et al. 2014; Aran et al. 2013), so we decided not to consider more distant sites to reduce false positives.

Figure 1 shows that there are hundreds to thousands of CpGs associated with each gene. The selection of important CpGs for each gene can be cast as a linear regression variable selection problem. The lasso is a commonly used variable selection technique that utilizes a L1-norm penalty on the regression coefficients to provide a sparse solution (Tibshirani 1996). The elastic net (Zou and Hastie 2005) is another alternative that utilizes a mixture of L1 and L2 penalty and works well when many of the predictors are correlated, which tends to be the case for methylation of nearby CpG sites (Leek et al. 2010). Both the lasso and the elastic net consider all predictors equally in variable selection, while it is known that certain CpGs sites are *a priori* more likely to be functionally important than others depending on the CpG type and relative location to the gene body, as described in Introduction. To incorporate such prior knowledge, we propose a sequential, two-step variable selection approach. Our sequential lasso method implements lasso regression in two steps, first considering CpG sites that are most likely to be associated with gene expression *a priori*, which are determined by a preliminary genome-wide study, and then checking the rest of the CpG sites to see if any of those can further explain the variability in gene expression.

### Notation and model

Let *y_ik_* denote the gene expression data for individual *i* (*i* = 1,…, *n*) at gene *k* (*k* = 1,…, *K*). Suppose that for gene *k*, there are *J_k_* CpG probes located within the gene body or in the flanking ±500kb on either end. Let *x_ijk_* denote the methylation beta value for individual *i* at CpG site *j* (*j* = 1,…, *J_k_*) of gene *k*. A standard linear regression model with independent and identically distributed Gaussian errors is assumed to relate gene expression with methylation:

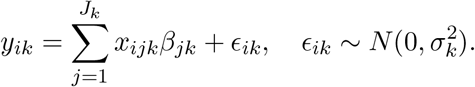

Or equivalently, in matrix form:

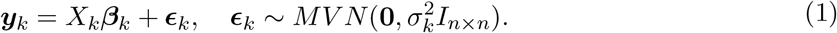

For gene *k*, our sequential lasso method performs variable selection among the *J_k_* associated CpG probes to identify a sparse subset whose methylation can best predict expression. For these selected CpGs, the regression coefficients *β_jk_* can be interpreted as weights that indicate the direction (positive or negative) and strength (absolute magnitude) of the association between methylation at CpG site *j* and expression of gene *k*.

Let *z*_*jk*_ = (*z*_*jk*1_,*z*_*jk*2_, *z*_*jk*3_, *z*_*jk*4_, *z*_*jk*5_ denote epigenomic characteristics of CpG site *j* associated with gene *k*, defined as follows:

1. *z*_*jk*1_ is a discrete variable denoting the location of the CpG site *j* relative to its associated gene *k*, i.e., whether it is upstream, downstream, or within the gene body.
2. *z*_*jk*2_ is a continuous variable denoting the distance of the CpG site *j* relative to its associated gene *k*, which is defined as follows:

a. If the CpG site is upstream of the gene’s TSS, *z*_*jk*2_ is distance from the TSS;
b. If the CpG site is downstream of the gene’s TES, *z*_*jk*2_ is distance from the TES;
c. If the CpG site is inside the gene, *z*_*jk*2_ is its relative distance from TSS to TES, with *z*_*jk*2_ = 0 for TSS and *z*_*jk*2_ = 1 for TES.
3. z_jk3_ is a discrete variable coding the type of the CpG site j relative to CpG island, i.e., whether it is located in a CpG island, a CpG shore, a CpG shelf, or an open sea.
4. *z*_*jk*4_ is a discrete variable coding the size of the gene *k*, i.e., whether gene *k* is a small gene (0 – 20kb), medium gene (20 – 80kb) or large gene (80 – 500kb).
5. *z*_*jk*5_ is a discrete variable coding the marginal direction of association between the CpG site *j* and the associated gene *k*. Let *ρ_jk_* denote the Pearson correlation between *y_k_* = (*y*_1*k*_,…, *y*_*nk*_)′ and *x_jk_* = (*x*_1*jk*_,…, *x*_*njk*_)′. A CpG site *j* is considered positively associated with expression of gene *k* if *ρ_jk_* > 0 and negatively associated if *ρ_jk_* < 0.

### Procedures

#### Step 1

A preliminary whole-genome analysis is conducted to build a model to estimate the probability that a given CpG site is found to be associated with expression based on its epigenomic characteristics *z_jk_*.

We accomplish this by first applying simple lasso regression of the expression data on the methylation data of CpGs located within the gene body or 500kb on either end for each gene *k* to get an estimate 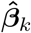, then modeling the probability of selection by lasso as a function of *z_jk_* (Wood 2006), i.e., Pr(*β_jk_* ≠ 0) = *f* (*z_jk_*). More specifically, for a CpG site *j* associated with gene *k*, its probability of selection by lasso is modeled as a smooth function of its relative location to the gene *k*, separately for each CpG location (upstream/downstream/inside gene), type, gene size category, and whether methylation of this CpG is marginally positively or negatively correlated with the target gene’s expression.

#### Step 2

For each gene *k*, lasso regression is applied to the methylation data of a subset of CpG sites that are *a priori* most likely to be associated with gene expression based on our model in Step 1.

A CpG *j* is considered *a priori* likely if *f* (*z_jk_*) > *π* for some threshold value *π*. Separate thresholds *π*_+_ and *π*_-_ are chosen for CpGs that are marginally positively or negatively associated with expression. Let the set 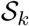 index the CpG sites *s.t. f* (*z_jk_*) > *π*_+_ for *ρ_jk_* > 0 and *f* (*z_jk_*) > *π*_-_ for *ρ_jk_* < 0. As seen in the results, the curves *f* have peaks at certain locations, and a tail that decays with distance from the gene body to a flat tail region. Here, the threshold values are chosen such that 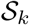 includes the CpGs located near the major peaks and excludes the CpGs from the relatively flat tails with low probability. Sensitivity analyses showed that our results were not overly sensitive to the exact choice of threshold if this general principle is followed.

Next, apply lasso regression to the expression data and the methylation data of the selected CpG sites in 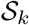, i.e.,

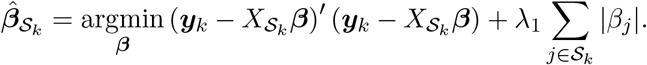

The regularization parameter λ_1_ is chosen by 5-fold cross validation. Denote the indices of the CpG sites with nonzero estimated coefficients among 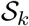 by 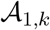. Regress ***y***_*k*_ on the methylation data in 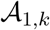 and get the residual vector *e_k_*.

#### Step 3

For each gene *k*, lasso regression is applied to methylation data of the rest of the CpG sites not considered in Step 2.

Treating the residuals *e_k_* from Step 2 as the new response, apply lasso regression to ***e**_k_* and the methylation data of the CpG sites in 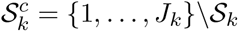, i.e.,

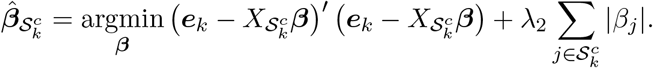

The regularization parameter λ_2_ is once again chosen by 5-fold cross validation. Denote the indices of the CpG sites with nonzero estimated coefficients among 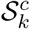 by 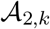.

#### Step 4

For each gene *k*, the regression model (1) is refitted by ridge regression only for the CpGs selected in Step 2 and 3.

Apply ridge regression to expression data ***y**_k_* and the methylation data of CpG sites in 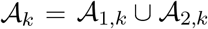, i.e.,

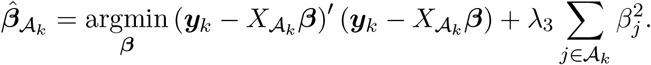

Again, the regularization parameter λ_3_ is chosen by 5-fold cross validation. Refitting the coefficients of CpG sites selected in Step 2 and 3 by ridge regression helps reduce the bias in estimation of large coefficients that plagues lasso regression (Zou 2006; Friedman et al. 2010).

This sequential lasso procedure gives priority to the CpG sites with characteristics that make them more likely to be selected as important based on information learned from the whole-genome analysis by letting them be selected first, and then other CpGs with characteristics that make them less likely to be important are only considered in the second step. This has several key benefits. First, it tends to lead to a parsimonious model that uses as few CpGs as possible in explaining the expression variability. Second, if multiple CpGs are correlated with each other and with expression, our approach will tend to select the one that is *a priori* more likely to be important (in 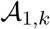) based on its characteristics rather than just arbitrarily selecting one, as a single-step lasso would do, which may make them more likely to be functionally important. Together, these contribute to a model that is more likely to discover biologically interpretable and reproducible results.

### Gene-specific methylation scores

Given GSMP, gene-specific methylation scores can be computed for each subject that can serve as gene-level summary of methylation and be used in graphical displays of methylation data or in integrative models to link different platforms at the gene level. GSMS gives a summary of the degree of functionally relevant methylation for a specific gene for a given subject.

For gene *k*, we rescale the predicted expression from the regression model 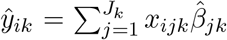 for subject *i* (*i* = 1,…,*n*), using the sample mean 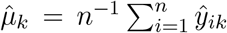 and standard deviation 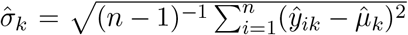, to compute the methylation score 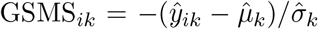. This rescaling puts the scores on a *Z*-score scale, which is intuitive and interpretable to most researchers in identifying subjects with unusually high and low levels of relevant methylation for this gene. The negative sign means that higher values of GSMS correspond to lower gene expression. In settings where the dominant signal is promoter methylation silencing the gene, this allows high GSMS to correspond to high promoter methylation and gene silencing. However, given that the relationship between CpG methylation and expression is not always this simple, care should be taken in interpreting these scores. Generally, high GSMS means that the methylation profile for that subject predicts a decrease in expression of the corresponding genes, while low GSMS means the methylation profile predicts higher gene expression.

## Data access

The TCGA data are publicly available. The gene expression and DNA methylation data of the MDACC Integromics cohort used for independent validation of GSMPs in CRC are provided in deidentified form in `Integromics.Rda`. The Shiny app and related R scripts are provided in `GSMP_Shiny.zip` which contain in R data files all of the GSMP results for the three tumor types in this paper, and will be expanded in the future to include other TCGA tumor types. These data and code will be deposited on the corresponding author’s GitHub repository and uploaded into appropriate public repositories upon acceptance of this paper.

## Acknowledgements

This work has been supported by CA-178744, CA-207101, CA-221707, CA-016672, NSF 1550088, the University of Texas M.D. Anderson Cancer Center Colorectal Cancer Moonshot, and the Del and Dennis McCarthy Distinguished Professorship Endowment in Gastrointestinal Cancer Research.

